# Aerosolized Desert Dusts Collected Near a Drying Lake and Pulmonary Inflammation in Mice: Implications for Environmental Exposures and Asthma

**DOI:** 10.1101/2022.09.07.506987

**Authors:** Trevor A. Biddle, Keziyah Yisrael, Ryan Drover, Qi Li, Mia R. Maltz, Talyssa M. Topacio, Jasmine Yu, Diana Del Castillo, Daniel Gonzales, Hannah M. Freund, Mark P. Swenson, Malia L. Shapiro, Jon K. Botthoff, Emma Aronson, David R. Cocker, David D. Lo

## Abstract

**Background:** A high incidence of asthma is prevalent among residents near the Salton Sea, a large inland terminal lake in southern California. This arid region has high levels of ambient particulate matter (PM); yet while high PM levels are often associated with asthma in many environments, it is possible that the rapidly retreating lake may contribute components with a specific role in promoting asthma symptoms.

**Objectives:** Our hypothesis is that asthma may be higher in residents closest to the Salton Sea due to chronic exposures to playa dust. Playa emissions may be concentrating dissolved material from the lake, with microbial components capable of inducing pulmonary innate immune responses. Such inflammatory responses may contribute to the development of asthma-like symptoms in residents. To test this hypothesis, we used a mouse model of aerosol exposures to assess the effects of playa dust.

**Methods:** From dust collected around the Salton Sea region, aqueous extracts were used to generate aerosols, which were injected into an environmental chamber for mouse exposure studies. We compared the effects of exposure to Salton Sea aerosols, as well as to known immunostimulatory reference materials. Acute 48-hour and chronic 7-day exposures were compared, with lungs analyzed for inflammatory cell recruitment and gene expression.

**Results:** Dust from sites nearest to the Salton Sea triggered lung neutrophil inflammation that was stronger at 48-hours but reduced at 7-days. This acute inflammatory profile and kinetics resembled the response to innate immune ligands LTA and LPS while distinct from the classic allergic response to *Alternaria*.

**Conclusion:** Lung inflammatory responses to Salton Sea dusts are similar to acute innate immune responses, raising the possibility that microbial components are entrained in the dust, promoting inflammation. This effect highlights the health risks at drying terminal lakes from inflammatory components in dust emissions from exposed lakebed.

## Introduction

The Salton Sea, a 345 mi^2^ body of water located in California’s Coachella Valley and Imperial Valley, is a site of frequent high levels of dust. Already designated a nonattainment area for particulate matter between 2.5 μm – 10 μm (PM10) and particulate matter under 2.5 μm (PM2.5) (California EPA Green Book 2022),^1^ dust in the region may soon worsen. Due to increasing temperatures and ongoing drought conditions, along with a 2003 ordinance shifting water away from the Sea, it has been estimated that 105,000 acres of additional playa will be exposed between 2003-2045 (Amato T. Evan 2019),^2^ with the shoreline having retreated substantially between 1985-2020 (Fig.1) Exposed playa has been linked to higher levels of dust in general (Reheis, 1997; Reynolds et al., 2007; Bullard et al., 2008; Hossein et al., 2018),^3–6^ and, in particular, the dust in the Salton Sea Basin is estimated to increase by 11% between 2018-2030^7^ (Parajuli and Zender, 2018) with changes in composition due to playa emissions (Frie et al., 2017).^8^

**Figure 1:**
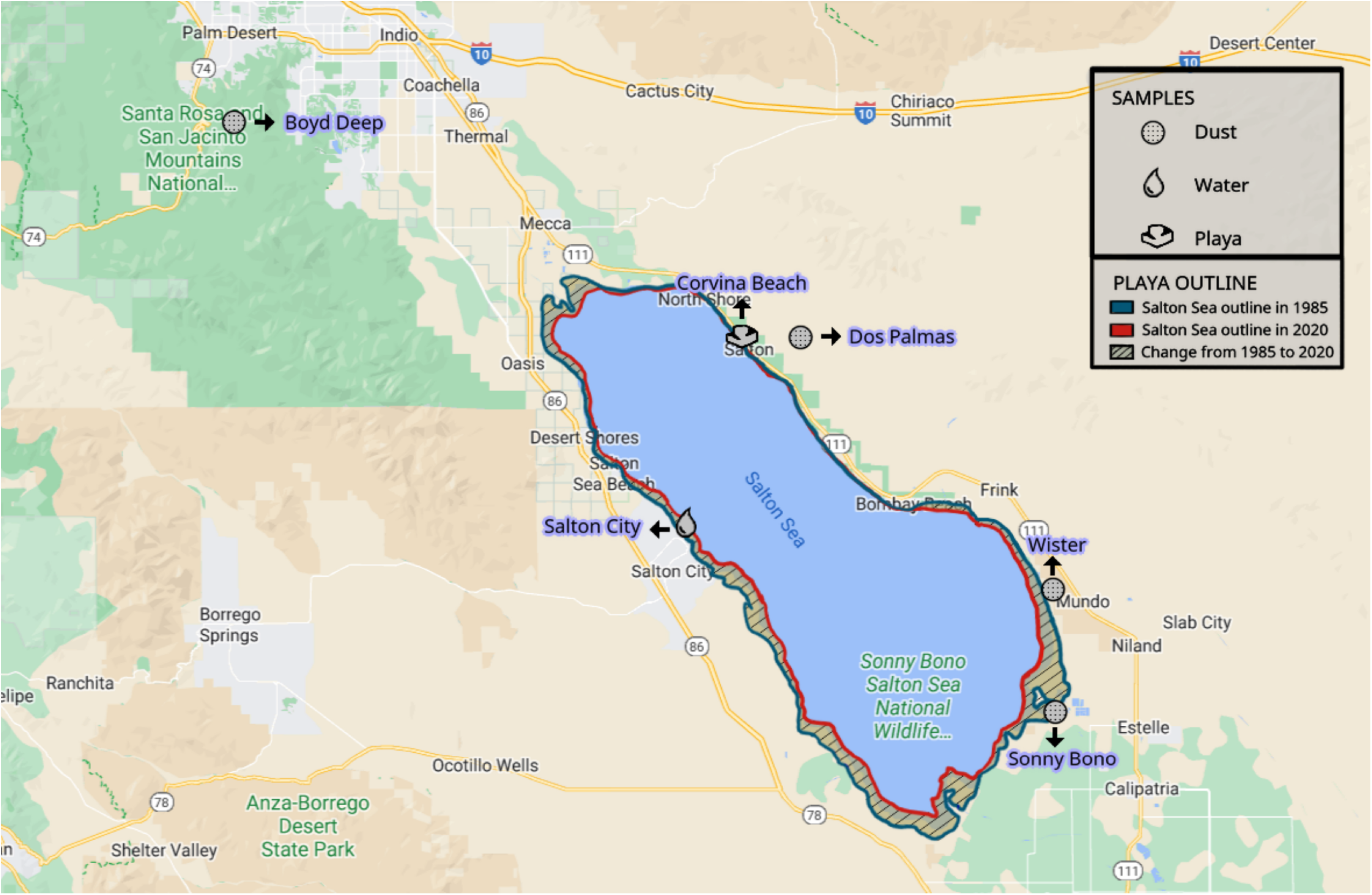
Collection Sites and Changes in Exposed Playa Over Time. Map showing the 3 dust collection sites (Wister, Dos Palmas, Sonny Bono) from around the Salton Sea, along with the desert dust control collection site (Boyd Deep Canyon), playa collection site (Corvina Beach), and water collection site from Biddle et al., 2021 (Salton City). Map also shows the change in playa exposure between 1985-2020 due to evaporation of the Salton Sea.

Along with high levels of dust, the communities surrounding the Salton Sea have high levels of childhood asthma. Currently, the asthma rate among children is estimated at 20%-22.4%, which is among the highest in the state of California, and noticeably higher than the state average of 14.5% (Farzan et al., 2019).^9^ Additionally, the communities surrounding the Salton Sea are among the top percentile for emergency room visits due to asthma (Calenviroscreen 4.0),^10^ though whether this is due to more severe asthma, lack of access to healthcare, high asthma rates, or a combination thereof is currently unknown.

Worryingly little is known about how the dust in the Salton Sea Basin may contribute to asthma and pulmonary disease. Asthma is generally defined as a disease of airway restriction, with an increase in airway hyperreactivity due to an allergen, along with immunoglobulin E (IgE) production, Th2 cytokine secretion and eosinophil recruitment (Bousquet et al., 2000).^11^ Other pulmonary diseases such as acute lung injury (ALI) instead preferentially recruit neutrophils (Blazquez-Prieto et al., 2018).^12^ Large dust storms, such as those in East Asia, have been linked to increased prevalence of respiratory diseases such as asthma (Chen et al., 2004; Kwon, Cho, Chun, Lagarde, & Pershagen, 2002; Watanabe et al., 2011),^13–15^ while organic dust from enclosed swine facilities has been shown to cause neutrophilic inflammation and oxidative stress (Poole et al., 2012; McGovern et al., 2016).^16–17^ Thus, dust can cause a variety of pulmonary disease; -however, it is unclear how Salton Sea dust may be contributing, especially in a location which serves as a reservoir for agricultural runoff. Contaminants such as pesticides, herbicides, heavy metals, along with microbial toxins, have been found in the Salton Sea (Carmichael and Li, 2006; Xu et al., 2016; Xu et al, 2016; Zhou et al., 2017).^18–20^ We previously showed that aerosolized Salton Sea water is capable of inducing gene expression changes indicating a mild inflammatory genotype without overt inflammatory cell recruitment (Biddle et al., 2021).^21^ However, in that paper we had not examined the potential contributions of dust in pulmonary disease.

While there have been previous studies examining the toxic effects of dust from the Salton Sea Basin, these studies relied on a single collected site and either an *in vitro* model (D’Evelyn et al., 2021),^22^ or intranasal administration of whole dust without a control dust site (Burr et al., 2021).^23^ As our study used an environmental exposure chamber to examine the potentially harmful effects of dust from multiple sites, including a desert dust control site, using an *in vivo* model while mimicking natural inhalation, we believe it offers a more robust and accurate look at dust as a source of pulmonary inflammation around the Salton Sea.

To understand the type of pulmonary inflammation caused by Salton Sea dust extracts, we used the aforementioned environmental exposure chamber, flow cytometry, and whole lung tissue gene expression. The inflammatory cell recruitment to the airways and lung tissue, along with changes in gene expression, were analyzed at either a 48-hour or 7-day timepoint and compared to well established acute, innate inflammatory agents LTA, LPS and the chronic, adaptive inflammatory allergen *Alternaria* to determine where the dust extracts fall along these axes.

## Methods

### Sample Collection

At each of four sites, we collected dust using passive collectors; our collectors consisted of a modified round bundt pan (Nordic Ware, Minneapolis, MN, USA) coated with Teflon (25.4 cm in diameter), which was lined with Kevlar mesh (Industrial Netting, Maple Grove, MN, USA), as described in Aciego et al. (2017)^24^ and Maltz et al. (2022).^25^ Glass marbles were suspended from the bottom and rested atop the mesh within the pan. Pans were fitted with overarching cross-braced strapping, which was covered in bird repellent (Bird-X, Elmhurst, IL, USA) to deter bird visitation or roosting. All pans, marbles, and mesh were acid washed in 2 M HCl, with rinses of 18.2 M Ω water between each reagent cleaning step, prior to deployment or contact with any dust. We deployed each of our collectors atop wooden posts, 2 m above ground level, within open canopy locations at each field site; this minimized the contribution of local dust inputs from nearby vegetation and local saltation.

On August 14th, 2020, we deployed these passive dust collectors at the “Boyd Deep” site, located in the Boyd Deep Canyon Reserve of the University of California (20 miles from the northwest border of the Salton Sea shoreline; see Supplemental Table 1). On August 30th, 2020, dust collectors were deployed at the “Sonny Bono” site, Sonny Bono Salton Sea National Wildlife Refuge, the “Wister” site, in the Wister Unit of the Imperial Wildlife Area, and the “Dos Palmas” site, in the Dos Palmas Preserve of the Bureau of Land Management. There was a second deployment at the “Wister” site starting on September 18^th^, 2021. These three sites are located at distances of 0.6, 2.0, and 4.0 miles from the nearest border of the Salton Sea lakebed, respectively (Supplemental Table 1; Fig. 1). After 81 days (for 2021 Wister dust), 55 days (for Boyd Deep Canyon dust), or 41 days (for Sonny Bono, 2020 Wister, Dos Palmas dust) these dust collectors were taken down, sealed in sterile Whirl-pak collection bags (Nasco, Madison, WI, USA), and transported to the University of California Riverside for further processing.

To sample playa, we used sterilized putty knives and flat spatulas to scrape the top layer (3.0 - 5.0 mm) of playa from the muddy exposed surface at 1.0 m - 2.0 m distances from the Salton Sea shoreline at Corvina Beach. From a 0.25 m^2^ plot, we slid large flat sterilized spatulas into playa material to sever the top layer of playa from the underlying sediment. We used the sterile putty knife to scrape this playa material into sterile Whirl-pak collection bags. Playa samples were frozen prior to subsequent transport and processing at the University of California Riverside.

### Sample Processing

To recover and archive dust samples from collectors across each of these date ranges, we used 18.2 M Ω water to extract the dust. We rinsed the marbles, mesh, and inner pan from each collector, using sterilized gloves to dislodge dust from these surfaces into the water suspension; we then removed the mesh and marbles and transferred the remaining water and dust suspension to acid-washed 1 L bottles (Nalgene Nunc International Corporation, Rochester, NY, USA; high density polyethylene; HDPE), for further processing and subsequent analyses.

All materials used for filtration and storage of samples were sterilized by acid washing, as described above. Frozen wet playa samples were weighed to 100 g and suspended into 1 L of 18.2 M Ω water within a beaker. Playa suspensions were mixed at the lowest speed on a stir plate at ambient temperatures and then filtered through autoclaved cheese cloth. We then filtered our playa suspensions into glass funnels through autoclaved 5.0 μm filters (Millipore-Sigma, Burlington, MA, USA).

We filtered all playa and dust suspensions into glass funnels through sterile 0.2 μm filters (47-mm diameter; Pall Supor 200 Sterile Grid filters, Pall Corporation, Port Washington, New York, USA) into collecting flasks. The remaining flow-through of playa (i.e., playa filtrate) was analyzed and subsequently used for mouse chamber exposures. Dust filtrates were frozen at a 45 ° angle in Fast Freeze Flasks, prior to lyophilization on a Labconco FreeZone 2.5 L −50 °C benchtop freeze dryer (i.e., lyophilizer, Labconco Corp., Kansas City, MO, USA). Lyophilized dust filtrate was subsequently normalized by dust mass (mg) in an aqueous suspension for use in mouse chamber exposures.

### Animals

All animal studies were done following the UCR institutional IACUC and NIH guidelines. Both male and female 8–9-week-old C57BL/6J mice were purchased from Jackson Labs, Sacramento and acclimated for one week in the University of California, Riverside SPF vivarium. After acclimation, mice were placed in an environmental exposure chamber for experimentation. Mice were kept 3-4 to a cage and allowed food and water *ad libitum*. A 12-hour day/night cycle was provided.

### Chamber operation

Exposure studies were performed in dual 540 L animal chambers (an exposure chamber and a control chamber) developed as described in Peng et al. (2019)^26^ and as used in Biddle et al. (2021).^21^ The relative humidity, temperature, and atmospheric pressure were measured in both chambers, with ammonia selectively measured in some exposures to ensure consistent exposure conditions were maintained. Mice in the exposure chamber were continuously fed a mixture of dry filtered air (0.5–1 lpm) and aerosolized spray (dried by two in-line silica gel columns, 3.5–4.5 lpm). The PM was generated from solutions of *Alternaria alternata* and *Alternaria tenuis* filtrate (Greer Laboratories, Lenoir, NC, USA; 0.4 g/L), lipoteichoic acid (LTA) from *Staphylococcus aureus* (Sigma Aldrich, St. Louis, USA), Lipopolysaccharide (LPS) from *Escherichia coli* O55:B5 (Sigma Aldrich, St. Louis, USA), and playa or dust collected from various sites in the region near the Salton Sea (0.025-0.100 g/L). The aerosol sources for the exposure were each tested in advance in order to determine chemical composition (high-resolution time-of-flight aerosol mass spectrometer (HR-ToF-AMS), Aerodyne) and aerosol density (aerosol particle mass analyzer (APM), Kanomax) to prepare the solutions to yield the targeted particle mass concentration. Sample aerosolization was accomplished by using a homemade nebulizer with silica-gel dryers (Peng et al., 2019).^26^ Mice in the control chamber were given filtered dry air (5.0 lpm) only, with other conditions matching the exposure chamber, including bedding replacement, food and water supplies, and corresponding day/night cycle. Particulate matter was only monitored within the exposure chamber by a scanning mobility particle sizer (SMPS, including Series 3080 Electrostatic Classifier and Ultrafine Condensation Particle Counter 3776, TSI) to assist in maintaining stable PM concentration. The target PM concentrations were as follows: 750 μg/m^3^ for the *Alternaria* mixture, 150 μg/m^3^ for LTA, 1 μg/m^3^ for LPS, and 1500 μg/m^3^ for environmental samples. *Alternaria*, LTA, and LPS doses were set at levels that promoted inflammatory cell recruitment without an accompanying cytokine storm (data not shown). The environmental sample concentration was similar to our previous studies in Biddle et al. (2021),^21^ where 1500 μg m^-3^ of water from Salton City was capable of inducing differential gene expression in the lungs by 7-days of exposure. For each exposure (n = 6-12), we used an equal number of male and female mice. Each exposure had a control air cohort that matched the number and sex of the exposure group. Mice were kept in the chamber for either 48-hours or 7-days depending on the study.

### Animal Processing

After either 48-hour or 7-days, the mice were removed from the environmental exposure chamber. They were then anesthetized using isoflurane and euthanized by cervical dislocation. Bronchoalveolar lavage fluid (BALF) was collected by flushing the lungs 3 times with 0.8 mL of PBS. Afterwards, the lungs were dissected out for digestion or RNA extraction. The right lung lobes were flash frozen in liquid nitrogen and kept at --80°C until RNA extraction. The left lobe minced into small (~1-2 mm) sections and digested using 0.5 mg/mL collagenase D (Roche Diagnostics, Mannheim, Germany) and 50 U/mL DNase I (Sigma Aldrich, St. Louis, USA) in RPMI 1640 (Gibco, Grand Island, USA) fortified with 10% heat-inactivated FB (Gibco, Grand Island, USA) preheated to 37°C. After incubating 30 minutes at 150 rpm in 37°C, the lung was agitated using an 18-gauge needle and incubate for another 15 minutes under the same conditions. Following digestion, the lung was pushed through a 100 μm cell strainer (Corning, Corning, USA). The cell strainer was then washed with RPMI 1640 with 10% heat-inactivated FBS before centrifugation and resuspension for use in flow cytometry.

### Flow Cytometry

BALF and post-digested lungs were centrifuged at 1500 rpm before resuspension in 100 μL of a 1:100 dilution of Zombie Yellow™ dye (Biolegend, San Diego, USA). After staining, cells were washed in FACS Buffer, centrifuged, and resuspended in 100 μL of a 1:50 dilution of Mouse BD FC block (BD Pharmigen, San Joe, USA). Cells were then stained using the following fluorescent antibodies: anti-CD45 FITC (BioLegend, San Diego, USA; Clone 30-F11), anti-CD19 Percp-Cy5.5 (eBioscience, San Diego, USA; Clone eBio1D3), anti-CD3 Alexa Fluor 700 (BioLegend, San Diego, USA; Clone 17A2) or anti-CD3 APC-Cy7 (BioLegend, San Diego, USA; Clone 17A2), anti-Ly6G BV510 (BioLegend, San Diego, USA; Clone 1A8), anti-CD11b BV421 (BioLegend, San Diego, USA; Clone M1/70), anti-CD11c PE-Cy7 (BioLegend, San Diego, USA; Clone N418) and anti-SiglecF APC (BioLegend, San Diego, USA; Clone S17007L).

Cells stained with Zombie Yellow^™^ dye were excluded from further analysis. Cell populations were determined using the following surface markers: neutrophils were CD45^+^CD11 b^+^Ly6G^+^SiglecF^-^CD11 c^-^, eosinophils were CD45^+^CD11 b^+^SiglecF^-^CD11 c^-^, T cells were CD45^+^CD3^+^SiglecF^-^CD11 c^-^, and B cells were CD45^+^CD19^+^SiglecF^+^CD11 c^-^

Samples were run on a NovoCyte Quanteon (Agilent Technologies, Santa Clara, USA). Gating and analysis were performed using FlowJo (Version 10.81, Ashland, USA).

### RNA Extraction

RNA was purified from the frozen right lung lobes using a TRIzol^©^ (Ambion, Carlsbad, USA) based method. ~100 mg of frozen lung tissue (half of the right lobe) was placed in a mortar, frozen with liquid nitrogen, and ground into dust using a pestle. After, the ground lung tissue was placed in TRIzol©. Chloroform was added, and the solution was mixed and centrifuged. The surface aqueous phase was mixed with isopropyl alcohol and centrifuged. The RNA pellet was then washed 3 times with 75% ethanol before drying at room temperature. This pellet was then resuspended in DEPC-Treated water (Ambion, Austin, USA). Concentration and purity of the RNA was checked via NanoDrop 2000 (Thermo Scientific, Carlsbad, USA).

### NanoString Analysis

50 ng of purified RNA was analyzed using an nCounter^®^ Sprint Profiler (NanoString Technologies, Seattle, USA) with the nCounter^®^ Mouse Immunology Panel according to manufacturer protocols. The nSolver^®^ 4.0 software (NanoString Technologies, Seattle, USA) was used to normalize gene counts based on housekeeping genes and positive controls. Differential expression was calculated using the nSolver^®^ Advanced Analysis 2.0 software (NanoString Technologies, Seattle, USA), and p-values were adjusted using the Benjamini-Hochberg method.

Normalized log2 gene counts and differential expression data were imported into R version 4.1.2^27^ (R version 4.1.2; R Core Team 2022) using *readr*^28^ (Wickham H et al., 2022) and *readxl*^29^ (Wickham and Bryan, 2022) and visualized using the *ggplot2*^30^ package (Wickham, 2016) and the *Khroma*^31^ package (Frerebeau N, 2022). Principal Component Analyses^32^ (PCA; Pileou 1984) were calculated using the normalized log2 gene count and the “prcomp” function.

This data was visualized using the *ggforitify*^33^ package (Tang et al., 2016). Dendrograms were generated using the average log2 gene count for each type of exposure and the built-in hclust function using the Ward.D2 method. They were visualized as an unrooted dendrogram using the *ape*^34^ package (Paradis E and Schliep K, 2019)

### Statistical Analysis

Statistical analysis for inflammatory cell infiltration was done using GraphPad Prism 9 (GraphPad, San Diego, USA). P-value was calculated using the Mann-Whitney U test for nonparametric data. Results shown include all mice, along with the average + the standard error (SE). P-value for gene expression was calculated using nSolver^®^ 4.0 and was false discovery rate (FDR) adjusted using the Benjamini-Hochberg method. A p-value of less than 0.05 and an FDR adjusted p-value of less than 0.1 were considered significant.

## Results

### Pulmonary Inflammation due to Salton Sea Dust Extract Compared to a Desert Dust Extract Control

To assess the biological effects of inhalation of these dusts in the Salton Sea region, we made aqueous extracts from collected dust, filtering out inert and larger particulate material. We injected suspensions of fine aerosols (~100nm diameter) into an environmental chamber for chronic exposures of mice. In previous studies on exposure to aerosols generated from Salton Sea water (“sea spray”), the exposed lungs showed induction of sets of genes associated with low level immune activation; however, no active tissue inflammation (i.e., recruitment of inflammatory cells such as neutrophils, eosinophils, lymphocytes) was detected above background (Biddle et al., 2021).^21^ In striking contrast, in mice exposed for 7 days to aerosolized extracts from Salton Sea dust collected at the Imperial Wister Unit, there was infiltration by granulocytes around the major airways (Fig.1b). Additionally, there was dramatic recruitment of neutrophils in bronchoalveolar lavage fluid (BALF) (13.9% + 8.8% vs 0.08 + 0.04%; Fig.1c). T cells were also preferentially recruited to the airways (2.4 + 0.7% vs 0.5 + 0.2%; Fig.1e), though B cells (Fig.1f) and eosinophils (Fig.1d) were not detected above background levels. Moreover, gene expression patterns were consistent with activation of acute immune inflammation (Fig.1h). The most highly expressed genes included several neutrophil chemokines (*Cxcl3, Cf2rb*), among other inflammatory chemokine (*Ccl6, Ccl9*). Additionally, several innate immune receptors (*Cd14, TLR2*) and inflammatory cytokine *Il1a* were upregulated in exposed mice.

For comparison, exposure to extracts of dust collected at Boyd Deep Canyon, a site in the desert distant from the prevailing winds of the Salton Sea, showed no significant inflammation in either the lung tissue (Fig.1a), BALF (Fig.1c,d,e,f), nor a gene expression pattern indicative of active inflammation (Fig.1g). While there were some inflammatory chemokines upregulated (*Cxcl3*, *Ccl9*), they were approximately 25-50% lower than in mice exposed to extract from around the Salton Sea. Additionally, control extracts failed to elicit upregulation in innate immune receptors or cytokines.

### Comparison of 48-Hour and 7-Day timepoints for Salton Sea Dust Extract and Reference microbial toxins LTA, LPS, and allergen Alternaria

Interestingly, although communities near the sea appear to suffer from a high incidence of asthma, the inflammatory response seen here did not show eosinophil recruitment nor Th2 gene regulation, which are hallmarks of allergic inflammation. This is not a limitation of the relatively short term (7d) aerosol exposure, as similar exposure to aerosols of the fungal allergen *Alternaria* produces robust allergic inflammation within 4 to 7 days (Peng et al., 2018; Biddle et al., 2021).^21,35^ Second, exposures of only 48 hours induced the strongest neutrophil recruitment (63.9 + 3.5% vs 0.1 + 0.03% in the airways; Fig2a; 41.2 + 10.4% vs 8.6 + 0.6% in the digested lung tissue; Fig.2c) with strong but slightly lower persistent inflammation present after 7 days of exposure (Fig.2a) and no recruitment of eosinophils at either timepoint (Fig.2b,d). Allergic stimuli such as *Alternaria* tend to have a stronger response at 7 days, as 48 hours does not appear to be enough to generate an adaptive response. Both neutrophilic and eosinophilic recruitment was greater at 7 days in *Alternaria* exposed mice (Fig.2a,b). Thus, material from dust at the Salton Sea induced strong pulmonary inflammation, but not the allergic profile more commonly associated with clinical asthma.

**Figure 2.**
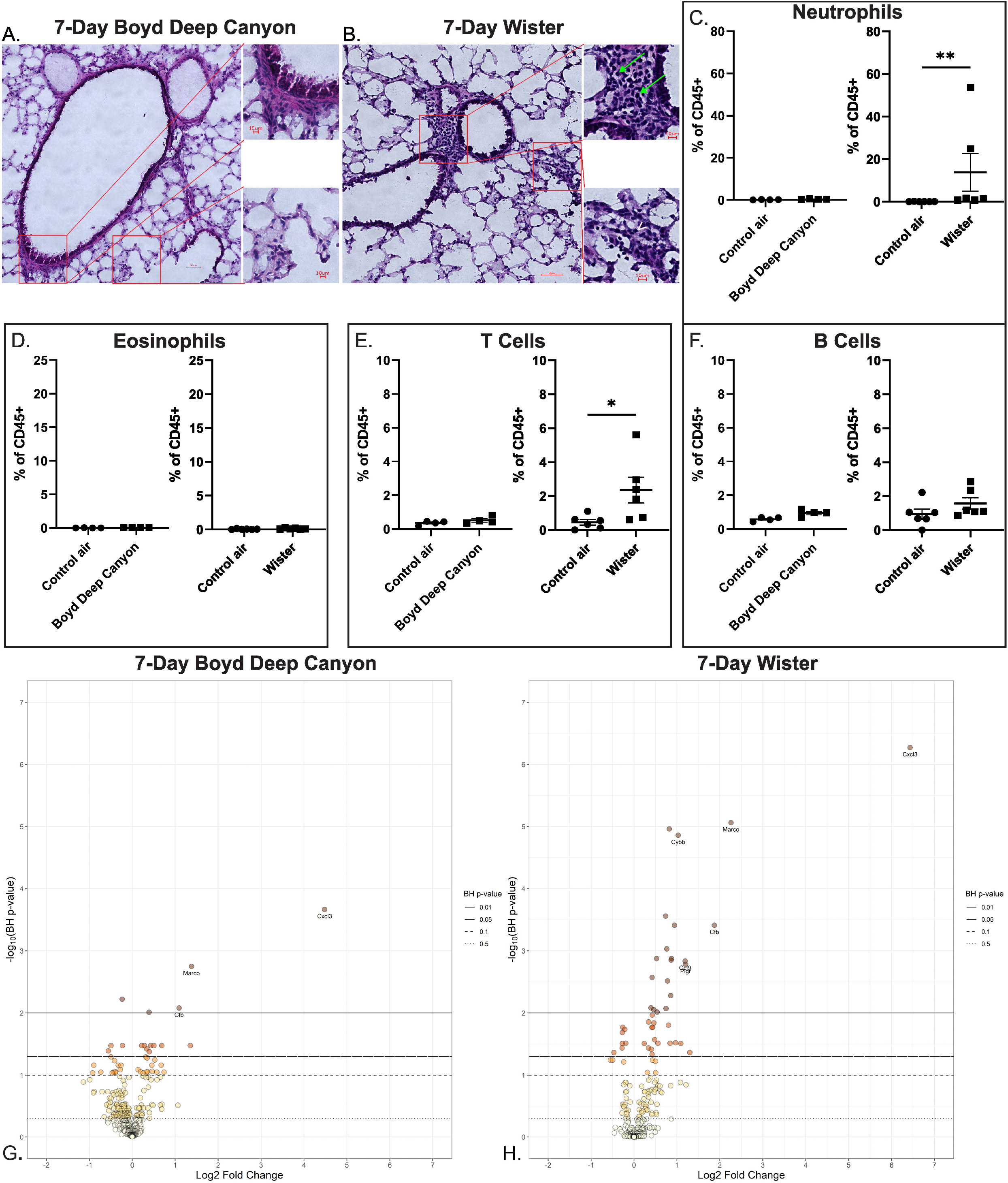
Pulmonary inflammation triggered by Salton Sea (Wister) Dust extract and by Desert (Boyd Deep Canyon) Dust extract. (A) 20X magnification H&E stain of lung tissue from mice exposed to Boyd Deep Canyon Dust extract. Insets are 60X magnification. (B) 20X magnifcation H&E stain of lung tisue from mice exposed to Wister Dust extract. Insets are 60X magnification. (C) Neutrophilic infiltration into the bronchoalveolar lavage fluid measured as a percentage of total immune cells (CD45+) as determined by flow cytometry. Neutrophil were defined as CD45^+^CD11b^+^Ly6G^+^SiglecF^-^CD11c^-^. (D) Eosinophilic infiltration into the bronchoalveolar lavage fluid measured as a percentage of total immune cells (CD45+) as determined by flow cytometry. Eosinophils were defined as CD45^+^CD11 b^+^SiglecF^+^CD11 c^-^. (E) T cell infiltration into the bronchoalveolar lavage fluid measured as a percentage of total immune cells (CD45+) as determined by flow cytometry. T cells were defined as CD45^+^CD3^+^SiglecF^-^CD11c^-^. (F) B cell infiltration into the bronchoalveolar lavage fluid measured as a percentage of total immune cells (CD45+) as determined by flow cytometry. B cells were defined as CD45^+^CD19^+^SiglecF^+^CD11 c^-^. All Boyd Deep Canyon Dust extract (*n = 4*) and Wister Dust extract (*n = 6*) exposed mice were compared to contemporaneous sex- and age-matched mice exposed to filtered house air. (G) Differential expression plot for Boyd Deep Canyon Dust extract exposed mice (*n = 6*) compared to contemporaneous sex- and age-matched mice exposed to filtered house air (*n = 6*). (F) Differential expression plot for Wister Dust extract exposed mice (*n = 9*) compared to contemporaneous sex- and age-matched mice exposed to filtered house air (*n = 9*). Gene expression was measured by a Nanostring Sprint Profiler using the mouse immunology panel; analysis was done using the accompanying nSolver software and visualized using *ggplot2*. Labeled genes have a log2 fold change greater than 1 or less than negative 1 and an Benjamini-Hochberg adjusted False Discovery Rate of < 0.01. * p < 0.05. ** p < 0.01. p-value determined using the Mann-Whitney U test for nonparametric data. Average + standard error shown.

The kinetics and types of inflammatory cells recruited matched conventional patterns of responses to innate immune triggers such as LPS and LTA, ligands for the innate receptors TLR4 and TLR2 respectively. These microbial components triggered strong, acute neutrophilic inflammation at 48-hours; however, over the course of chronic 7-day exposure the inflammatory response was significantly attenuated. The attenuation over the course of 7 days for Salton Sea dust extracts and innate immune triggers vs the increasing response after exposure to allergen is also shown at the gene expression level. The average log2 fold change decreased in all three exposures, from 0.476 to 0.219 for Wister extract, 0.266 to 0.177 for LPS, and 0.243 to 0.191 for LTA but increased from 0.226 to 0.683 for *Alternaria*. By the 7-day timepoint, the Salton Sea dust extracts, LPS, and LTA had attenuated to the point that their gene expression patterns clustered closer to the mice given only filtered air while the allergic inflammation gene expression patterns in mice exposed to Alternaria were clearly distinct at 7-days (Fig.2e). This tendency towards resolution is not seen in an allergic model, as the *Alternaria* response matured over the course of 7 days.

### Gene Expression Comparison of Multiple Salton Sea Collection Sites

To confirm that neutrophil recruitment and innate immune activation were representative of the entire Salton Sea basin rather than specifically the Imperial Wister Unit, we exposed mice to dust extract collect from two additional sites around the Salton Sea, the Dos Palmas Preserve and Sonny Bono. As the response was higher at 48-hours for dust extract from the Imperial Wister Unit, we used this timepoint for analysis. We found that dust extract from these locations triggered neutrophil infiltration to the BALF and gene expression profiles (Fig.S1) consistent with dust collected from the Imperial Wister Unit. This gene expression profile clustered with the other innate immune triggers and the Alternaria mixture before maturation into a Th2 response (Fig.3a). By contrast, mice exposed to extracts of playa (dry exposed lakebed), one of the major potential sources of dust, failed to recruit significant neutrophil responses and showed few gene expression changes (Fig.S2), with only Marco and Il1a being significantly upregulated.

**Figure 3.**
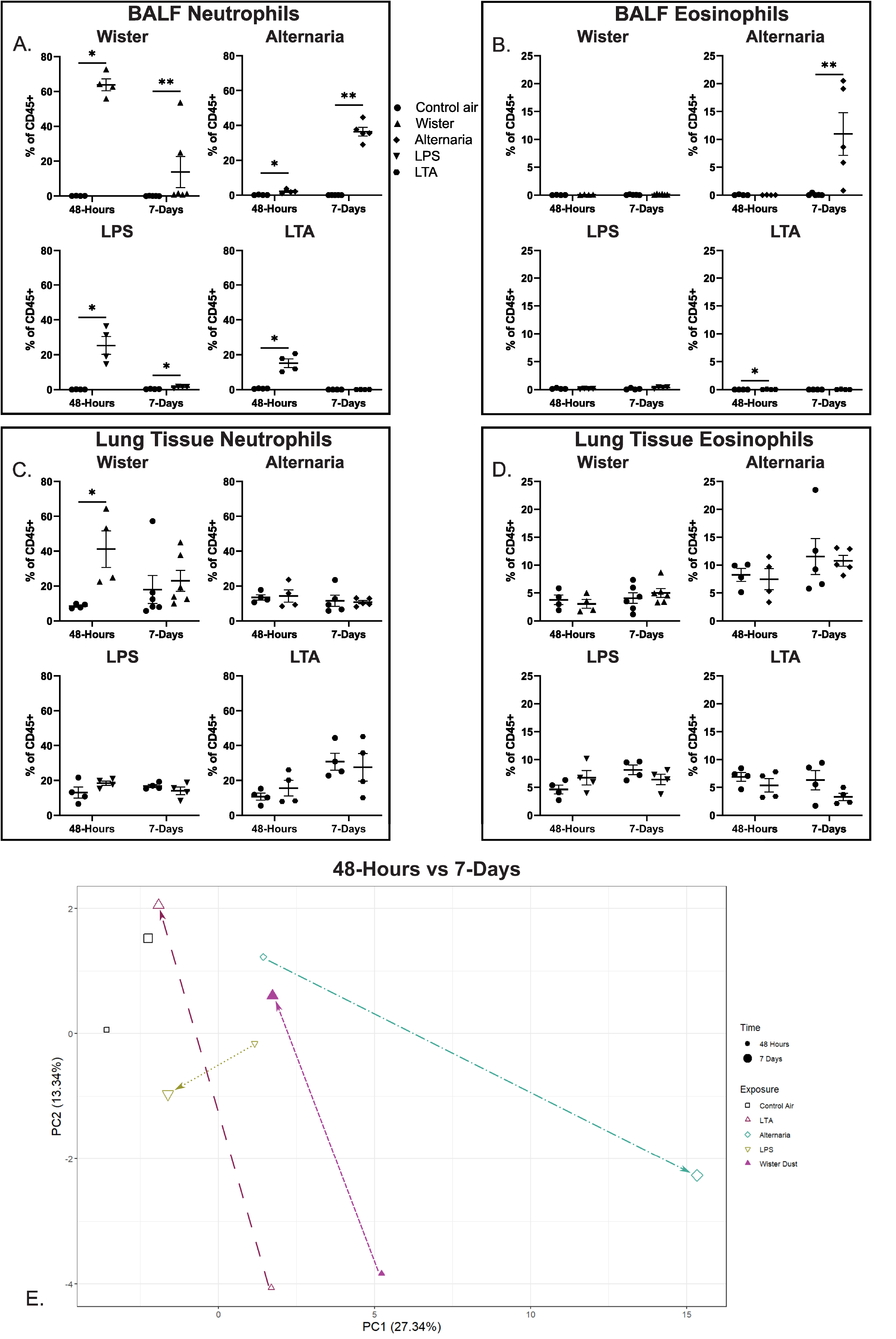
Changes from 48-hour to 7-day timepoints for Salton Sea (Wister) Dust extract, *Alternaria*, LTA, and LPS exposed mice. (A) Neutrophil infiltration at 48-hours and 7-days in the bronchoalveolar lavage fluid (BALF) as a percent of total immune cells (CD45+) for Wister Dust extract, *Alternaria*, LTA, and LPS exposed mice. (B) Eosinophil infiltration at 48-hours and 7-days in the bronchoalveolar lavage fluid (BALF) as a percent of total immune cells (CD45+) for Wister Dust extract, *Alternaria*, LTA, and LPS exposed mice. (C) Neutrophil infiltration at 48-hours and 7-days in the lung tissue as a percent of total immune cells (CD45+) for Wister Dust extract, *Alternaria*, LTA, and LPS exposed mice. (D) Eosinophil infiltration at 48-hours and 7-days in the lung tissue as a percent of total immune cells (CD45+) for Wister Dust extract, *Alternaria*, LTA, and LPS exposed mice. Neutrophils were defined as CD45^+^CD11b^+^Ly6G^+^SiglecF^-^CD11c^-^, while eosinophils were defined as CD45^+^CD11 b^+^SiglecF^+^CD11 c^-^. All timepoint and exposure combinations were compared to contemporaneous sex- and age-matched mice exposed to filtered house air. *n = 4-6* (E) PCA plot showing the changes from the 48-hour timepoints to 7-day timepoints for Wister Dust extract, *Alternaria*, LTA, and LPS exposed mice. PCA was generated using the *prcomp* function in R and visualized using *ggplot2*. Each point represents the average location of the specific timepoint and exposure combination on PC1 and PC2. Arrows highlight trend line from 48-hour to 7-day for each exposure. *n = 6-9*. * p < 0.05, ** p < 0.01. p-value calculated using the Mann-Whitney U test for nonparametric data. Average + standard error shown.

To better characterize the relationship between the chronic exposures, we performed a similar analysis using our 7-day exposures. We found that there was significant overlap between the control air and the Boyd Deep Canyon Dust extract exposed mice. LTA exposed mice, whom had the most significant attenuation of inflammation by 7-days, also overlapped heavily with the control air. The LPS and Wister Dust extract exposed mice were distinct from the controls, but not to the extent of the *Alternaria* exposed mice. Thus, while LPS and Wister Dust do not have a maturation of the immune response by 7-days, they appear to have not fully resolved, indicating a potential for low-level inflammation over a longer time course (Fig.3b). It should be noted that LPS and Wister responses were distinct from each other, with minimal overlap. Thus, they are unlikely to be triggering inflammation through the same mechanism.

The relationship between the different exposures is summarized in the dendrograms in Fig.3c and Fig.3d. For the 48-hour exposures, the response to playa is at one end, with the majority of exposures on the other branch. The Wister dust has the longest branch, as it has the strongest response and is unique among the other exposures. The others are closely related, with minor separation. For the 7-day exposures, Alternaria is entirely separated from the other exposures. The control dust extract is on the opposite side from the innate immune-like responses, which all have little distinction from each other.

## Discussion

The response to Salton Sea dust showed similarities to innate immune ligands LTA and LPS in the overall kinetics and patterns of gene induction. These responses started with high levels of neutrophil, but not eosinophil, recruitment to the airways before decreasing to nonexistent (LTA) or minimal (LPS) levels by 7 days of continuous exposure, along with an attenuation of inflammatory gene expression. This is wholly distinct from an adaptive immune Th2 response to the allergen Alternaria which started with a low level of neutrophil recruitment before maturing into a stronger response highlighted by high levels of neutrophils, eosinophils, and an increase in genes related to Th2 responses. As the Salton Sea dust followed the kinetics of the innate immune ligands, it appears unlikely to be triggering an adaptive response.

Yet the effects of Salton Sea dust did not strictly match reference innate ligands. Salton Sea dust triggered neutrophil recruitment that was persistent for at least 7 days, which was not the case in mice exposed to LTA. Additionally, we found upregulation of *TLR2* in our Salton Sea Dust exposed mice, but not in LTA or LPS exposed mice, indicating potential for exacerbation after dust exposure, even in the case of selective tolerance to the dust. All Salton Sea dusts induced persistent upregulation of *Il1a* through 7 days of exposure; no other 7-day exposure in our study induced significant upregulation of *Il1a*. *Il1a* has been associated with neutrophil recruitment by cigarette smoke (Botelho et al., 2011),^36^ as well as promoting inflammatory cytokine release and inhibiting fibrotic ECM release and healing by lung fibroblasts (Suwara et al., 2013; Osei et al., 2016).^37–38^ The persistent upregulation of *Illa* could be a key culprit in the pattern of inflammation caused by chronic Salton Sea dust exposure and contribute to long term lung pathology without conforming to either microbial toxin-driven innate nor adaptive immune patterns of immunity. This is important as other innate immune triggers such as LPS tend to attenuate and provide immune tolerance (Natarjan et al., 2010),^39^ which may not be the case in Salton Sea dust exposure.

It is important to clarify that the patterns of inflammation compared in this study are similar to familiar innate immune triggers composed of microbial components (cell wall, endotoxin, etc.) that are not by themselves directly toxic, and are mediated instead through innate immune receptors such as TLR2 and TLR4. These should be viewed as rather distinct from the effects of actual microbial toxins (e.g., cyanotoxins, microbicidins, etc.), produced by a variety of microbes including both bacteria and algae. The effects of these types of toxins on lung inflammation are not within the scope of this study, and while Salton Sea water is known to have detectable levels of some cyanotoxins (Carmichael and Li, 2006),^18^ it is not known if the cyanotoxins can aerosolize and cause pulmonary inflammation nor it is known whether Salton Sea dust contains harmful concentrations of these toxins. However, studies have cited the potential impact of microbial toxins in triggering or exacerbating asthma symptoms when present in red tides (Fleming et al., 2007; Zaias et al., 2011).^40–41^

In addition to the ways that Salton Sea dust may directly cause pulmonary disease, the dust may interact with allergens to modify asthma development. Exposure to LPS during allergic sensitization to ovalbumin or house dust mite can worsen Th2 asthma or shift it towards a corticosteroid resistant Th17-type asthma (Li et al., 1998; Barboza et al., 2013; Yu and Chen, 2018; Sadakane et al., 2018; Thakur et al., 2019).^42–46^ As Salton Sea dust triggers inflammation similar to LPS, it may work in similar ways. This would have significant implications for the communities surrounding the Salton Sea, as different forms of asthma require different treatments.

The similarity of the Salton Sea dust responses to innate immunity that can be triggered by “Pathogen-Associated Molecular Patterns” – that is, bacterial components such as zendotoxin, cell wall, and other materials – raises the question of whether specific microbial species in the Salton Sea or accompanying playa dust are particularly pro-inflammatory in lungs. While a comprehensive microbiome analysis of the aeolian microbiota has yet to be published, several studies have analyzed the composition of the Salton Sea and the sediment. Proteobacteria, particularly Gammaproteobacteria and Alphaproteobacteria, make up the majority of the bacteria in the Salton Sea and in the sediment, followed by Bacteroidetes (Dillon et al., 2009; Swan et al., 2010; Hawley et al., 2014).^47–49^ These types of bacteria are known major contributors to LPS in the gut (d’Hennezel et al., 2018),^50^ and can be carried on dust particles over large distances (Tang et al., 2017),^51^ indicating a potential source of microbial inflammatory substances, particularly LPS. Understanding the interplay between the water, playa and aeolian microbiota is likely critical to understanding the pulmonary inflammation due to Salton Sea Dust (Freund et al., 2022).^52^

As our preparation method filters out whole bacteria, our results would be due to the products produced by the bacteria rather than an immune response towards the bacteria itself. Additionally, while our results were similar to LPS induced pulmonary inflammation, it was not a perfect match. Thus, while LPS may be a contributing factor to dust-driven pulmonary inflammation, it is unlikely to be the only factor.

In sum, our results provide insight into the pulmonary health effects of a drying terminal lake. The dust collected from around the Salton Sea was uniquely toxic when compared to desert dust collected from a location protected from the prevailing winds around the sea. While there is likely a unique ecology and contaminants to the Salton Sea Basin, we believe the more general features of the drying lake, associated aerosol dusts, and consequent health impact, has broad relevance to other regions plagued by chronic drought and drying lakes. These regions are unique as they have rapidly increasing levels of dust, which has the unique property of triggering high levels of pulmonary inflammation that is likely to worsen the pulmonary health of already vulnerable nearby communities. By understanding the inflammation triggered by the dust, the mechanisms behind the inflammation, and how it interacts with allergens and allergic development, we may be better equipped to handle the negative health impacts of the dust and developed tailored strategies to address them.

**Figure 4.**
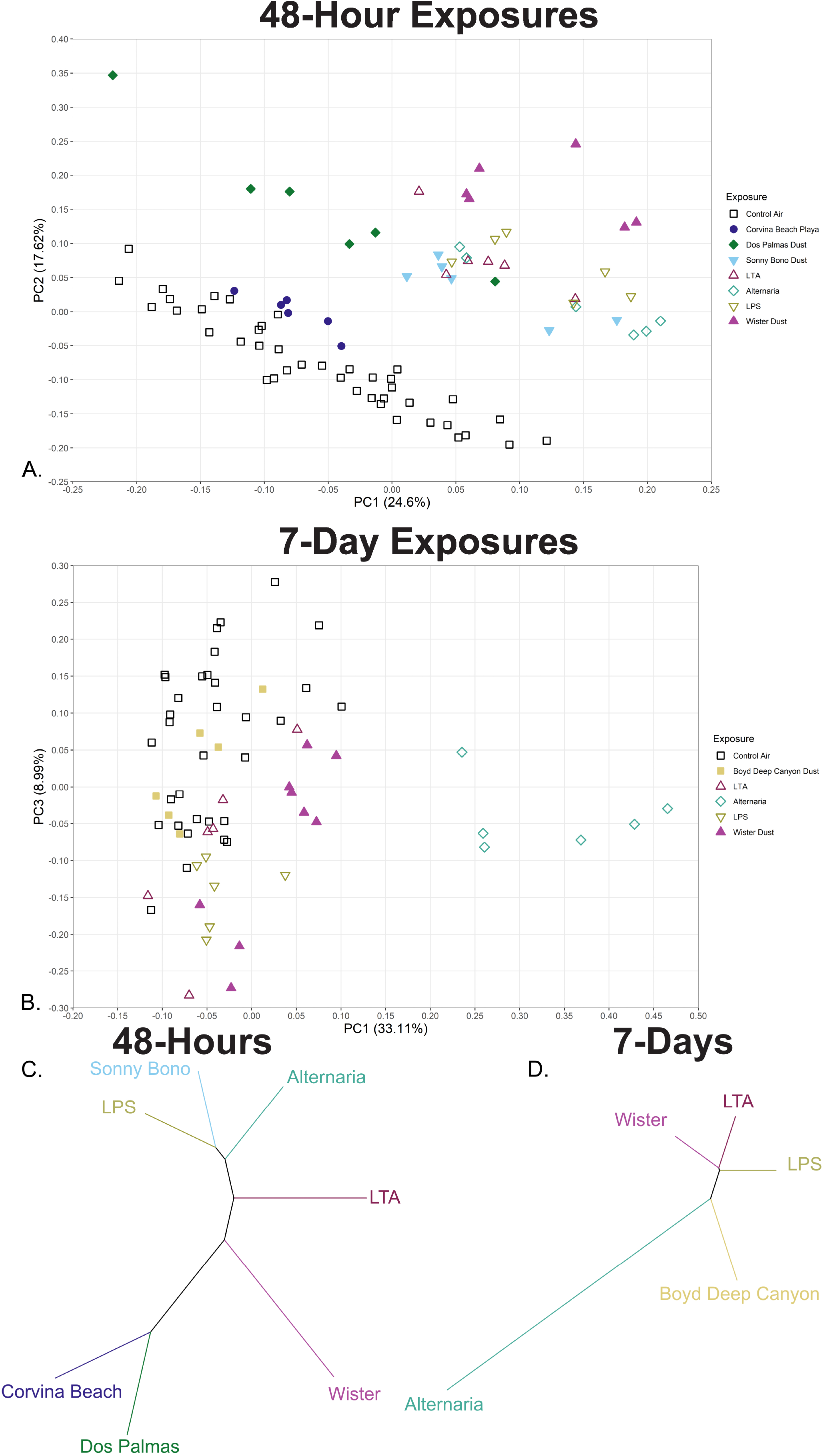
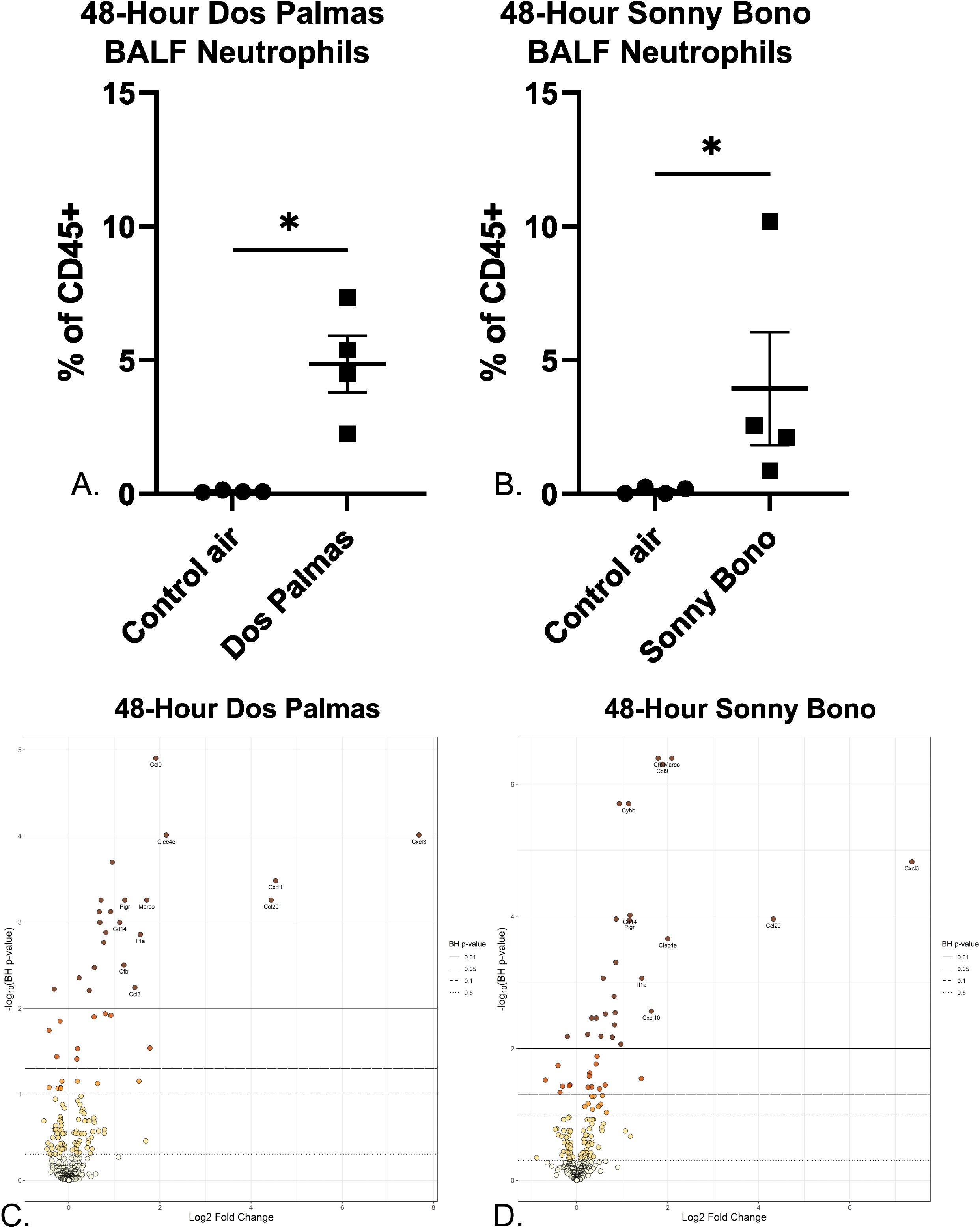
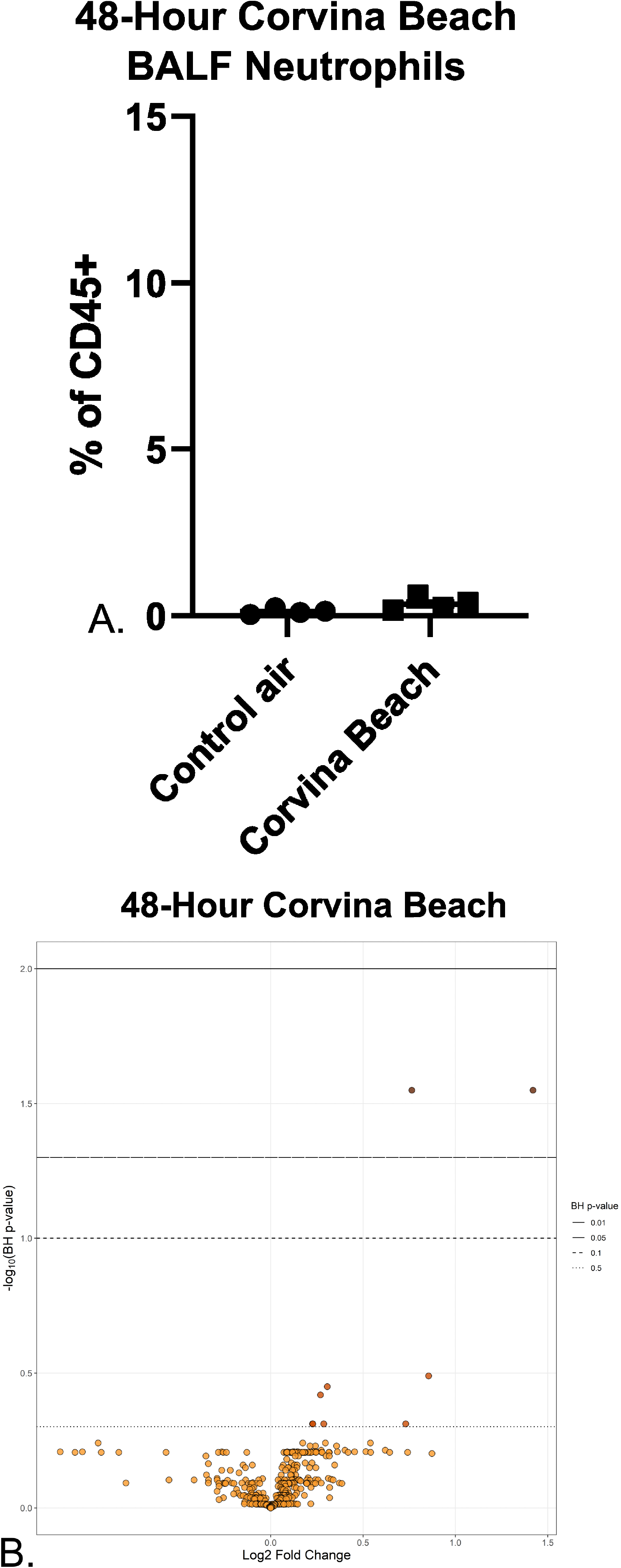
Gene expression comparisons for 48 hour and 7-day timepoints. (A) PCA plot comparing Salton Sea Dust extract (Wister, Sonny Bono, Dos Palmas), Salton Sea Playa extract (Corvina Beach), LTA, LPS and *Alternaria* exposed mice at 48-hour (*n=6*). (B) PCA plot comparing Salton Sea Dust extract (Wister), Desert Dust extract (Boyd Deep Canyon), LTA, LPS and *Alternaria* exposed mice at 7-days (*n = 6-9*). PCA was generated using the *prcomp* function in R and visualized using *ggplot2*. (C) Dendrogram showing the relationship between Salton Sea Dust extract (Wister, Sonny Bono, Dos Palmas), Salton Sea Playa extract (Corvina Beach), LTA, LPS and *Alternaria* exposed mice at 48-hour (*n = 6*). (D) Dendrogram showing the relationship between Salton Sea Dust extract (Wister), Desert Dust extract (Boyd Deep Canyon), LTA, LPS and *Alternaria* exposed mice at 7-days (*n = 6-9*). Dendrograms were generated using the log2 value for each gene averaged by the exposure. This average was then used with the hclust function (method = Ward.D2) and visualized using the *ape* package.

## Acknowledgments

The research presented in this publication was supported by the National Institute On Minority Health And Health Disparities of the National Institutes of Health under Award Number U54MD013368 to DDL. The content is solely the responsibility of the authors and does not necessarily represent the official views of the National Institutes of Health. TAB lead experiments and was the primary author of the manuscript. KY, TMT, DDC and MLS helped with mouse dissections and processing. RD, QL, and DG ran and monitored environmental exposure chamber. HF, MPS, MRM collected and processed dust. JY assisted in creation of figure 1 JKB designed environmental dust collectors. EA and DRCIII helped with experimental design and processing. DDL assisted with experimental design and editing the manuscript.

## Notes

### Competing Interest Statement

The authors have declared no competing interest.

